# Age- and caste-independent piRNAs in the germline and miRNA profiles linked to caste and fecundity in the ant *Temnothorax rugatulus*

**DOI:** 10.1101/2023.05.21.541611

**Authors:** Ann-Sophie Seistrup, Marina Choppin, Shamitha Govind, Barbara Feldmeyer, Marion Kever, Emil Karaulanov, Alice Séguret, Sivarajan Karunanithi, Miguel V. Almeida, René F. Ketting, Susanne Foitzik

## Abstract

Social insects are models for phenotypic plasticity: the generation of different phenotypes from the same genotype. Ant queens and workers differ not only in their morphology and behaviour, but also in their fecundity and lifespan, which is often several times higher in queens. However, the gene regulatory mechanisms underlying these differences are not yet well understood. Since ant queens can live and reproduce for more than two decades, they need to protect their germline from the activity of transposable elements (TEs). This protection may be redundant in short-lived, often sterile workers. We have analysed the expression of two protective classes of smallRNAs, microRNAs (miRNAs) and Piwi-interacting RNAs (piRNAs), in different tissues, castes, and age classes of the ant species *Temnothorax rugatulus*. We show that piRNAs are particularly active in the ovaries of queens. TEs are clear targets of the piRNAs in this ant species, and piRNA-specific sequence signatures in the ovaries of all queens regardless of age indicate that young and old queens have similarly active piRNA pathways. Interestingly, the reduced ovaries of the workers also showed the same level of piRNA activity. This was not only the case in young, fertile workers from queenless nests, but also in the presumably older foragers, which have almost completely regressed ovaries. These findings suggest that the germline in these ants is invariably protected by piRNA activity, irrespective of ovarian development. The brain and thorax of queens also contained piRNAs, but at lower levels, and the piRNA-specific ping-pong signatures were strongly reduced in these tissues. We also annotated and analysed miRNAs in different tissues. We confidently detected the expression of 304 miRNAs. Of these, 10 were enriched in the brain and three to the thorax, whereas 83 were specific to the ovaries. 105 miRNAs were found to be expressed in all three tissues. We also identified miRNAs whose expression potentially is related to ant caste, fecundity, and age, suggesting that caste-specific gene activity may be regulated in part by miRNAs. In contrast, our studies of piRNA activity indicate similar profiles across caste, fecundity and age groups, but strong tissue specificity with the highest piRNA mediated TE protection in the germline.

## Introduction

The origin of eusociality is considered one of the major transitions in evolution (Szathmáry & Smith, 1995). Insect societies show a high level of complexity; their colonies have even been described as superorganisms (Boomsma & Gawne, 2018; Wheeler, 1911) in which the queen functions as the germline and the workers represent the soma. In ant colonies, for example, queens and workers are closely adapted to their specific roles. Ant queens focus on reproductive activity and are exceptionally long-lived, particularly in monogynous societies, i.e. societies with only a single queen (Keller & Genoud, 1997), wherein some queen ants can live for more than 20 years. In contrast, the sterile workers show a high behavioural complexity and take over all other tasks in the society, from foraging to brood care. Their lifespan is considerably shorter; they often live for less than one year. The different castes of social insects have been long regarded as model systems of phenotypic plasticity, as they are largely not genetically determined but arise due to differential transcriptional activity in the larval phase (Corona et ak,, 2016). While a strong investment in reproduction shortens the lifespan in many organisms (Flatt, 2011; Maklakov & Chapman, 2019), social insect queens can maintain - especially in comparison to workers – a high fertility and long lifespan. Indeed, a high fecundity has been shown to be positively linked to a long lifespan in ants, even within a single caste (Heinze & Schrempf, 2012; Negroni et al., 2020).

Our previous studies investigated the molecular regulation of ageing and fecundity in the small ant *Temnothorax rugatulus*, in which queens can reach lifespans of two decades. We saw that an increase in queen fecundity can result in an upregulation of longevity pathways such as DNA repair and autophagy (Negroni et al. 2021). Moreover, workers that developed their ovaries upon queen removal showed an extended lifespan and transcriptional shifts in TOR and insulin-like/IGF-1 signalling (Choppin et al., 2021; Negroni et al., 2021; Negroni et al., 2020). Gene expression changes with age in ant queens as well, with middle-aged queens investing more strongly in antioxidants, but less in immunity and starvation resistance, compared to young queens (Negroni et al., 2019). Although we have gained insights into the molecular pathways of ageing and fecundity in this species and its different castes, we still know little about the regulatory mechanisms underlying these transcriptional shifts. All that has been shown is that histone acetyltransferase activity plays a role in the molecular regulation of ovarian development in workers in the absence of the queen (Choppin et al., 2021). We also do not know how the strong transcriptional differences between the castes, which lead to these divergent phenotypes, arise and are maintained.

In many social hymenopteran species, worker ants may develop their ovaries and lay male-destined haploid eggs, a phenomenon known as arrhenotokous worker reproduction (Bourke, 1988). This occurs in response to specific social or environmental conditions, in particular when the queen dies or is lost. Indeed, the development and expression of reproductive traits in worker ants are tightly regulated by social and environmental cues, which ensure the maintenance of the reproductive division of labor within the colony. In our model species, the ant *T. rugatulus*, workers start fighting over dominance and develop their ovaries within weeks of queen loss. These workers do not only show activated ovaries but also an extended lifespan and altered regulation of mTOR or insulin-like / IGF-1 signaling (Choppin et al., 2021; Negroni et al., 2021; Negroni et al., 2020).

In this study, we aimed to investigate and describe small regulatory RNAs (sRNAs) from the ant *Temnothorax rugatulus* and address the question of whether sRNA signatures are tissue-specific and whether they may change with the age and reproductive status of the queen and worker castes. Different types of sRNAs exist, but they all act in concordance with a protein of the Argonaute protein family and typically they repress gene acvtivity (Ender & Meister, 2010). Two of the major sRNA pathways are the microRNA (miRNA) pathway (reviewed in Bartel, 2009) and the Piwi-interacting RNA (piRNA) pathway (reviewed in Ozata et al., 2019). In this study, we focussed on the expression of these two types of small RNA.

miRNAs are transcribed from distinct loci and the resulting transcripts form hairpin structures. These hairpins are processed by specific nucleases to produce mature miRNAs (Bartel, 2009). After binding their respective Argonaute proteins (AGOs), miRNAs direct the binding of this AGO protein to specific mRNAs, based on base-pairing between the miRNA and the mRNA, and repress their activity (Filipowicz et al., 2008). As miRNAs do not bind their target mRNAs with 100% sequence complementarity, it can be difficult to predict which genes a certain miRNA may target, and many tools exist to try and tackle this task (Bartel, 2009; Lewis et al., 2003). The targets of miRNAs are typically protein coding genes that have regulatory roles themselves (Bartel, 2009).

Much like miRNAs, piRNAs are transcribed from distinct loci in the genome. However, unlike miRNAs, which can be found in any tissue, piRNAs are often restricted to the germline. This is at least true for model organisms like *Drosophila*, *Caenorhabditis elegans*, and mice. However, in various arthropods, somatic piRNAs have been detected (Lewis et al., 2018) suggesting that the ancestral state of piRNA expression may have been ubiquitous. The processing of piRNA precursor transcripts into mature piRNAs does not proceed via a hairpin intermediate and makes use of a different, complex molecular machinery that has been reviewed in detail in (Zhang et al., 2022). Mature piRNAs are bound by AGO proteins of a specific sub-family: that of the Piwi proteins. Piwi::piRNA complexes are known to elicit strong silencing effects, both at the transcriptional and post-transcriptional levels (Ozata et al., 2019). Of note, when a Piwi::piRNA binds a target, it may turn the targeted mRNA into a new piRNA in a process that is known as the ping-pong cycle (Luo & Lu, 2017; Ozata et al., 2019). This process results in sense and antisense piRNAs that overlap precisely 10 nucleotides at their 5’ ends. Finally, piRNAs have a strong bias towards uracil at the 5’-position, matched by an adenine at position 10, due to the decribed ping-pong cycle. A deeply conserved role of piRNAs is the silencing of transposable elements (TEs) and the protection from non-self DNA (TEs), (Madhani, 2013). As a consequence, piRNA sequences are typically not conserved, even between closely related species, as they co-evolve with the sequences of the TEs that they are trying to control.

## Materials and Methods

### Ant collection and maintenance

*Temnothorax rugatulus* ants are widespread in western North America and live in high-elevation coniferous forests, under rocks or in crevices. In August 2015 and 2018, we collected several hundred colonies at various locations in the Chiricahua Mountains (see Choppin et al., 2021; Negroni et al., 2019 for GPS information on exact locations). Collection permits for the Coronado National Forest were obtained through the Southwestern Research Station of the Museum of Natural History in Portal, Arizona. The ant colonies were transported to our laboratory in Mainz, Germany, where they were relocated to individual boxes with plastered flours containing nests made of plastic inserts between two glass plates. The nests were covered with red foils to block the light. The colonies were kept in climate chambers at 18°C and 70 % humidity in a 12:12 light-dark cycle. They were fed with honey and crickets and received ad libitum water.

### Analysis of miRNA and piRNA expression depending on queen age and tissue

This project aimed to investigate miRNA and piRNA expression in different tissues of old and young ants queens. We were interested in whether only the ovarian tissue, i.e. the germline itself, uses piwi protection, or whether this is also found in other tissues, in this case the brain or fat body. Furthermore, we wanted to investigate whether this protection against TEs is lost in old queens or maintained over their entire lifespan. Seven young queens were collected during our 2018 collection trip as founding queens either alone or with their first few workers (always less than six workers at the time of sampling). These queens were probably less than 6 months old and established their colony in July or August of that year. Seven old queens were collected in Arizona in August 2015 in large monogynous colonies, indicating that, at the time of collection, they were likely several years old. We kept them for another 3 ½ years so that at the time of sampling, we estimate their age to be between 6-12 years. *Temnothorax* ant queens can live for up to 20 years (Plateaux, 1986). At the time of sampling, colonies of these young and old queens comprised between 60 and 230 workers. In November 2018, the queens were provided with fresh boxes, nests and food and housed in a climate chamber at 25°C and 70% humidity on a 12:12 light-dark cycle. All these large queens were from single-queen colonies.

After one month under the same conditions, the queens were dissected in ice-cold PBS under a Leica stereomicroscope. The head with the thorax, the fat body (first cuticular segment of the abdomen containing fat cells), and the ovaries were removed from each queen, separately frozen in liquid nitrogen, and stored at -80°C. We measured ovariole length and counted the number of white eggs (i.e., eggs in development) in the ovaries using the Leica software LAS v4.5. One young queen had died before the dissection date and was therefore not sampled. Differences between old and young queens was assessed using R. For the number of eggs, we used a a generalized linear model (glm(eggs ∼ age, family=quasipoisson)), and for the lengths, we used a linear model (lm(length ∼ age))

In August 2020, the brains of the queens were isolated from their heads on ice, placed in 50µl TRIzol, frozen in liquid nitrogen, and stored at -80°C. The brain of a young queen was not removed because her head was damaged. The brain was placed in 50µl TRIzol, frozen in liquid nitrogen, and also stored at -80°C. Two weeks later, the RNA was extracted from the brain and thorax samples. For this, 50µl chloroform was added to the samples, the tubes were swirled and placed on normal ice for five minutes. The samples were then centrifuged at 12000xg and 4°C for 15 minutes. The resulting upper phases were transferred to new 1.5mL Eppendorf tubes using gel loading tips. Then 25µL of 100% ethanol was added to the upper phases, the tubes were swirled and the samples were transferred to Zymo-Spin IC columns (Direct-zol RNA Microprep Kit from Zymo Research) in collection tubes. The collection tubes containing the columns were centrifuged at 12000xg for 30 seconds at room temperature, and the columns were then transferred to new collection tubes. 400µl RNA wash buffer was added to the columns, which were then centrifuged at 12000xg for 30 seconds at room temperature. 5µl DNase I was carefully mixed with 35µl DNA digestion buffer and added to the columns. The samples were incubated for 15 minutes at room temperature. Then 400µl Direct-zol PreWash was added to the columns and they were centrifuged at 12000xg for 30 seconds at room temperature. The residues were discarded, and the previous step was repeated. 700µl of RNA wash buffer was added to the columns, which were then centrifuged at 12000xg for one minute at room temperature. The columns were transferred to RNAse-free tubes and 15µl of DNAse/RNAse-free water was added directly to the column matrices. Finally, the collection tubes containing the columns were centrifuged at 12000xg for 30 seconds at room temperature to elute the RNA and the columns were discarded. The RNA samples were stored at -80°C until sequencing. In November 2020, 50µl of ice-cold TRIzol was added to the ovary samples and the RNA was extracted using the same procedure as described above.

### Analysis of miRNA and piRNA expression in ovarian tissue of queens and workers

Our second aim was to compare miRNA and piRNA expression in the ovaries of queens and workers of different fertility stages to investigate whether all females or only the queen protects her germline from the activity of transposable elements by piRNAs. In our focal species, *T. rugatulus*, the removal of the queen induces younger workers dedicated to the task of nursing to develop their ovaries and start laying eggs. We could demonstrate that the increase in fecundity of these brood carers was associated with an extended lifespan and altered immunity (Negroni et al., 2021; Negroni et al., 2020). In contrast, older workers, which serve as foragers, have often completely degenerated ovaries and are no longer able to reproduce. They could be regarded as the soma of the *superorganism* ant colony (Wheeler, 1911). Hence, colonies of *T. rugatulus* female ants display a gradient in terms of reproductive potential and life expectancy. Both traits are most pronounced in the queen. Only she can mate, and her ovaries also possess a spermatheca and eight ovarioles. The ovaries of the workers are smaller and are composed of only two ovarioles. Fertile nurses in ql colonies are the longest-lived and most fertile individuals after the queen. They are followed by the nurses from qr colonies (colonies with a queen) and finally by the old foragers from these colonies. This gradient allows us to analyse at which point these female ants stop protecting their germline from TEs or whether germline tissue always expresses this Piwi protection.

At the end of 2021, 12 qr and 12 ql colonies (colonies lacking a queen) with between 20-100 workers were provided with fresh boxes, nests and food and housed in a climate chamber at 25°C and 70% humidity on a 12:12 light:dark cycle. We aimed to sample four independent replicates of each of the four groups: queens, qr brood carers (qr-n) and foragers (qr-f) and ql nurses (ql-n). After one month, in January 2022, we dissected the ovaries of the queen, four brood carers and five foragers from 12 qr colonies each, and the ovaries of four brood carers from 12 ql colonies. For each queen replicate, the ovaries of three queens from three different colonies were pooled. For each brood carer replicate (qr and ql alike), we pooled the ovaries of four workers from three colonies, for a total of 12 ovaries. For the forager replicates, we pooled the ovaries of five foragers from three colonies, for a total of 15 ovaries each. Dissections were conducted in ice-cold PBS under a stereomicroscope and ovaries were photographed using the Leica system LAS v4.5. We noted the number of eggs in development and grouped the ovaries of queens and workers into five developmental stages (0 = regressed ovaries: thin, clear tubes; 1 = undeveloped ovaries with rounded tubes and a wider end, 2 = slightly developed ovaries containing immature eggs; 3 = developed ovaries with one mature egg, 4 = well-developed ovaries with 2-5 mature eggs, 5 = extremely well-developed ovaries with > 5 eggs). Dissections took less than 5 minutes and ovaries were transferred into 100µl Trizol, which was kept on ice until all ovaries of one pooled sample were added and then stored at -80°C. Differences between groups was assessed using R. For the number og eggs, we used a generalized linera mixed effects model (glmer(eggs ∼ group, family = poisson+ (1|Colony)) and for the developmental stages, we used a glm (glm(stage ∼ group, family = poisson)).

Each sample was ground while frozen and then RNA was extracted with the Zymodirect Zol Microprep following the standard instructions. RNA was resolved in 15µl RNAse/DNAse free water and stared at -80°C before sequencing.

### Sequencing

NGS library prep was performed with NEXTflex Small RNA-Seq Kit V3 following Step A to Step G of Bioo Scientific’s standard protocol (V19.01) using the NEXTFlex 3’ SR Adaptor and 5’ SR Adaptor (5’rApp/NNNNTGGAATTCTCGGGTGCCAAGG/3ddC/and5’ GUUCAGAGUUCUACAGUCCGACGAUCNNNN, respectively). Libraries were prepared with a starting amount of 5ng and amplified in 21 PCR cycles (first dataset, different tissues of old and young queens), or a starting amount of 1.5ng and amplified in 22 PCR cycles (second dataset; ovaries of different castes).

Amplified libraries were purified by running an 8% TBE gel and size-selected for 15 – 40nt. Libraries were profiled in a High Sensitivity DNA Chip on a 2100 Bioanalyzer (Agilent Technologies) and quantified using the Qubit dsDNA HS Assay Kit, in a Qubit 2.0 Fluorometer (Life Technologies). All samples were pooled in equimolar ratio and sequenced on 1 Highoutput NextSeq 500/550 Flowcell, SR for 1x 84 cycles plus 7 cycles for the index read.

### Read pre-processing, mapping, and filtering

NGS library quality was assessed using FastQC v0.11.8 (Andrews, 2010) before removing the constant sequence of the 3’-NextFlex adapter using Cutadapt v2.4 (Martin, 2011) (-a TGGAATTCTCGGGTGCCAAGG -m 26 -M 43). Trimmed libraries were once again assessed for quality using FastQC.

Reads were mapped to the draft genome assembly “trug_v1.0” (BioProject PRJNA750352, in submission) using Bowtie v1.2.2 (Langmead et al., 2009) allowing for 1 mismatch whilst concomitantly removing the random 2x 4nt random NextFlex adapter bases (bowtie -p 16 -v 1 -M 1 -y --best –strata --trim5 4 --trim3 4). Reads lengths were assessed by a custom Python script (https://github.com/Tunphie/SequencingTools/blob/main/summarizeNucleotideByReadLength.py) and plotted using ggplot2 (Wickham, 2016) in R v3.13. Mapped reads were subsequently split into different lengths for further analysis of piRNAs (reads of length 25-30nt) and miRNAs (reads of length 18-24nt) using samtools v1.9 (Danecek et al., 2021) and GNU awk. These mapped reads were further split into reads overlapping or not with miRNA or transposon annotation in sense or antisense orientation using the function intersectBed from bedtools v2.27 (Quinlan & Hall, 2010). This was done in order to determine lengths and 5’-biases using the aforementioned Python script before plotting these with ggplot2 in R.

### piRNA analyses

Reads in the piRNA-range (25-30nt) existing outside of our annotated transposons were split into plus and minus strand using samtools v1.9. BigWig files were then generated for each strand separately using the deepTools v3.5.1 (Ramírez et al., 2016) function bamCoverage. Means of groups were calculated using WiggleTools (Zerbino et al., 2014) and converted back into BigWig files using wigToBigWig from UCSC. computeMatrix and plotProfile functions from deepTools v3.5.1 were used to generate views of the mapped data coverage over selected loci, provided as a BED file.

Reads in the piRNA-range (25-30nt) overlapping with our annotated transposons (BioProject PRJNA750352, annotation in submission) were counted using Subread v2.0.0 (Liao, Smyth, & Shi, 2013)(featureCounts -s 1 -M -F SAF), thereby summarizing read counts per transposon of the same type (n-1903). Reads in different transposon families were subsequently summarized in R by comparison to the used annotation file. T-tests and subsequent Holm-correction for multiple testing were carried out in R. Differential analyses were carried out using DESeq2 v1.30.1 (Love et al., 2014) in R. A false discovery rate (FDR) or 0.05 was used. PCA plots and MA plots were generated using plotPCA and plotMA from the DESeq2 package.

Samples were randomly downsized to 500 thousand (old and young queens; thorax, brain, and ovaries) or 2 million (ovaries of queens, nurses, and foragers) reads per sample for our two datasets respectively using samtools collate and shuf (-n 500000 / -n 2000000). These reads were then used to search for ping-pong signatures using PingPongPro v1.0 (Uhrig & Klein, 2019)(-p -v -o). Ping-pong pairs were compared to the TE annotation and divided into signatures existing within TEs and outside of TEs using R. PingPongPro was further run using the TE annotation as input, and mapping relative to TE families was counted using R. Plots were generated using ggplot2 in R.

### miRNA prediction and annotation

We used miRDeep v.2.0.1.3 (Friedländer et al., 2012) on reads of length 18 to 24 nt to predict miRNA *de novo*. We first used mapper.pl script (-d -e -h -i -j -m -o 30 -v -p -s -t) to process the reads and map them to our partial *T. rugatulus* genome. The outputs from mapper.pl along with our partial genome and miRNA annotations from *A. mellifera*, *D. melanogaster*, *T. castaneum*, and *B. mori* were used to predict miRNAs using the miRDeep2.pl script (-g -1 –P). We quantified the predicted miRNAs using the quantifier.pl (-d) script. Further, we removed predicted miRNAs that did not satisfy the following quality filters adapted from (Coenen-Stass et al., 2018). The miRNAs should: i) have a miRDeep2 score above 1, ii) have a statistically significant miRDeep2 randfold p-value (p<0.05), iii) have a sequence that does not match any tRNAs or rRNAs in RFAM (Griffiths-Jones et al., 2003) and iv) have at least one read in three or more of our samples from our first dataset (old and young queens; thorax, brain, and ovaries). The parameters which were set in contrast to default settings are mentioned in parentheses following the respective scripts.

We do note that some of our reads may be misclassified; either as the wrong sRNA type or as a sRNA when it was indeed the breakdown product of a functional RNA. We noted that some piRNAs were more likely the breakdown product of rRNAs (Figure 3D&F-G and Figure S2C), and that some nonTE-derived piRNA reads more resembled structural reads (Figure S1D). Some piRNAs may also be misclassified as miRNAs due to the closeness of their length distributions and their similar 5’-bias.We do, however, believe that most piRNAs would be filtered out of our differential miRNA analyses since these are the only reads mapping sense to our miRNA annotation. We also note that sRNA populations other than miRNAs and piRNAs exist, and that some of our sRNAs may have been misclassified due to not investigating further sRNA families such as small interfering RNAs (siRNAs).

### miRNA analyses

Reads in the miRNA-range (18-24nt) aligning to our miRNA annotation were counted using Subread v2.0.0 (featureCounts -s 1 -M -F SAF --minOverlap 18). miRNA expression for a given tissue (brain, ovary, and thorax) was defined as non-zero read counts in at least three samples from that tissue regardless of age. Overlap of these reads were plotted using VennDiagram in R (Chen & Boutros, 2011). Differential analyses were carried out using DESeq2 v1.30.1 in R. A false discovery rate (FDR) of 0.05 was used. PCA plots and MA plots were generated using plotPCA and plotMA from the DESeq2 package.

## Results

### Basic sRNA profiles of three *T. rugatulus* tissues

In order to investigate the tissue-specific expression of sRNAs in *T. rugatulus*, we sequenced sRNAs shorter than 35 nucleotides (nt) in length from the thorax, ovaries, and brain of young, founding queens below six months of age and middle-aged established queens above 3.5 years. When comparing the general read length distribution, two distinct populations were observed; one in the typical miRNA range (18-24nt) and one in the typical piRNA range (25-30nt)(Figure 1A). The peak in the piRNA range was more prominent in the ovaries than in the somatic tissues, consistent with findings in other animals showing that the piRNA pathway is highly active in the germline. In contrast to germ cells, the miRNA-range peak was more prominent in the somatic tissues, although a distinct piRNA-range peak could also be observed in these tissues (Figure 1A). To probe deeper into the characteristics of the sRNA molecules, we aligned the reads to a draft genome assembly from Jongepier et al. (2021). Below, we will describe the miRNA- and piRNA-range sRNAs and soma and germline in separate sections. Lastly, the difference between old and young queens will be described.

**Figure 1:**
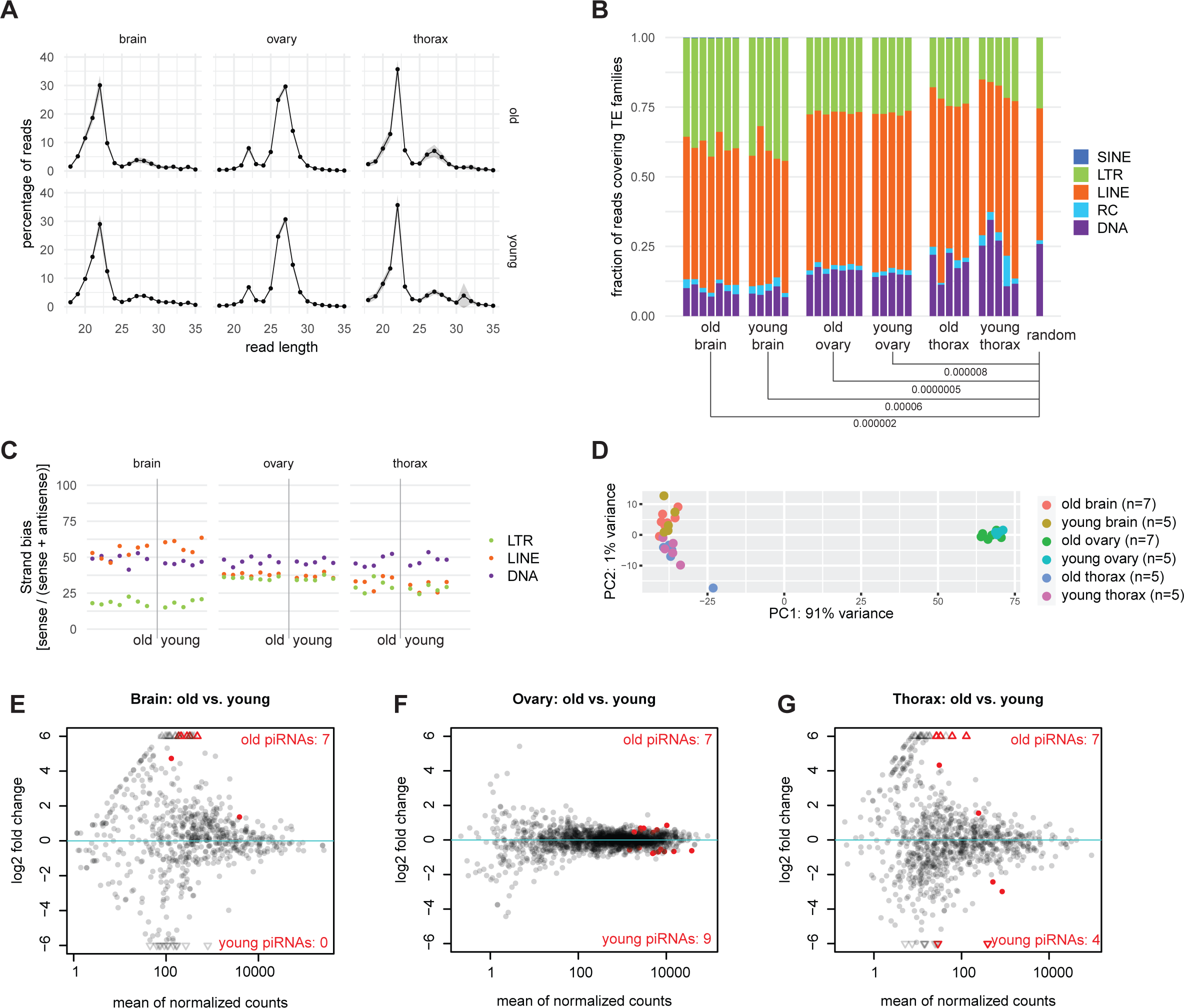
piRNA expression in different tissues of old and young queens. **A** Length distribution of all reads generated by sRNA-Seq. Shaded areas depict standard deviation, obtained from the 5-7 biological replicates. **B** Stacked bar plots showing piRNA-like reads covering different TE families. T-tests carried out for each TE family to find difference between old and young and between each group and annotation. For each pair of comparisons, the lowest Holm-adjusted p-value is shown, if this is significant (p<0.05). All non-significant values not shown but listed in the supplementary data. **C** Strand bias of piRNA-like reads derived from LTR, LINE, and DNA elements in the three tissues, obtained from young and aged queens. **D** Principal component analysis of piRNA-like reads (25-30nt) that map to annotated TEs. **E-G** MA plots showing the pairwise comparisons of piRNA targeting of specific TEs in old vs. young queens in the three tissues. Each dot represents one TE. Red indicates significant changes (FDR<0.05). All hits are reported in the extended data, Table 1.

### Queen ovaries express TE-targeting piRNAs

After having determined that ovaries have a prominent piRNA-like sRNA population which peaked at abundance at lengths of ∼26-28nt, we extracted all reads in the range 25-30nt and compared these to a TE annotation provided by Jongepier et al. (2021), since piRNAs are known to be involved in TE repression, particularly in gametes. We saw that many reads, up to 60%, overlapped with annotated TEs in the queen ovaries (Figure S1A). We further extracted these TE-derived reads and saw that they mapped to both sense and antisense strands of TEs, with some bias for the antisense strands, as has been previously shown for piRNAs in other species (Figure S1B). The reads furthermore had a strong bias for U at the 5’-position; another common feature of piRNAs (Figure S1B). Compared to the genome-wide TE annotation, DNA and Rolling-circle derived piRNAs were slightly underrepresented in the piRNA-like reads in the ovaries, whereas LTR- and LINE-derived reads were mildly overrepresented (Figure 1B).

In the absence of available gene annotation, we could not determine where the piRNA-like reads that were not assigned to TEs mapped. Nevertheless, the reads mapping outside of TEs also had a 5’-U bias and showed mapping on both genomic strands similar to the TE-derived sRNAs (Figure S1C). We did notice that some loci were strongly biased to produce sRNAs from only one of the strands (Figure S1D), suggesting that these may not be true piRNAs. Alternatively, these may represent highly strand-biased populations of piRNAs, as described before (Brennecke et al., 2007; Hirano et al., 2014). Due to the absence of a detailed genome annotation, this could not be analysed further

These results are consistent with the idea that this 25-30nt sRNA population corresponds to piRNAs. To further assess this, we probed for so-called ping-pong signatures. These arise when cleaved piRNA target transcripts are themselves turned into piRNAs. Such piRNAs will display a characteristic 10nt overlap on their 5’ end with the piRNA that induced the cleavage. We first downscaled all libraries to 500K reads in order to enable comparison between them, and then searched for possible ping-pong pairs using PingPongPro (Uhrig & Klein, 2019). We found around 4,000 high-fidelity ping-pong pairs in the ovaries across the entire genome, 67% of which were found inside annotated TEs (Figure S1E). We then looked for ping-pong pairs specifically in the TEs and noticed that the majority of them were derived from LTR and LINE elements with more than 50% of TE-derived ping-pong signatures stemming from LTR elements (Figure S1F). Given that LTR elements only amount to 25% of the annotated transposons, this represents a significant overrepresentation, and suggests that these TE types are most actively targeted by the Piwi pathway. In the absence of functional antibodies, which would allow us to test whether the identified sRNAs are bound to Piwi proteins, we conclude that the reported sRNA species represent piRNAs, and that *T. rugatulus* ovaries express an active piRNA pathway, with LTR retrotransposons as important targets.

### Thorax and brain have limited piRNA expression in queens

Next, we probed the thorax and the brain of queens for the existence of piRNAs. In both tissues, we did find a population of small RNAs that displayed similar features to the piRNAs we found in the ovaries with length peaks at roughly 27nt (Figure 1A). They showed a mild bias towards antisense polarity with respect to their matching transposons, and they displayed a strong bias for a 5’-U (Figure S1B). Interestingly, the brain had more piRNAs mapping outside of annotated TEs (Figure S1A&C-D), possibly pointing towards a function of piRNAs other than transposon surveillance in this tissue.

In contrast to ovaries, ping-pong signatures were practically absent in the brain (0-200 signatures per 500K reads per sample). Some high-fidelity ping-pong signatures, around 500 per 500K reads (overall average: 459, average in old samples: 572, average in young samples: 346), were found in the thorax (Figure S1E), but their abundance was still roughly 5-10-fold lower compared to ovaries. Overall ping-pong results were not strongly affected by age (Figure S1E).

Then, we checked how the available piRNAs were distributed over the different TE families. In the thorax, the piRNAs were distributed along the annotated transposons roughly according to their representation in the genome, albeit with high variance between samples (Figure 1B). Interestingly, the brain samples showed a consistent overrepresentation of LTR elements at the expense of DNA elements (Figure 1B). These results could point to differential activities of TE families in these two tissues.

Curiously, we noticed that the strand bias of LTR and LINE elements differed in the brain relative to the ovaries and thorax (Figure 1C): LTR-derived piRNAs showed a much more extreme anti-sense bias, whereas LINE-derived populations showed a weaker anti-sense bias and even a mild over-representation of sense piRNAs. These findings may also reflect differential TE activities in the brain compared to the thorax and ovaries.

### piRNA expression differs minimally between old and young queens

We saw no difference in TE family targeting or strand bias between old and young queens (Figure 1B&C). In order to further determine whether ageing of *T. rugatulus* queens is associated with differential targeting of specific TEs by piRNAs, we carried out differential analyses of this subset of sRNAs using DESeq2 (Love et al., 2014).

The principal component analysis revealed that the largest differences in piRNA expression existed between ovarian and somatic tissues, irrespective of age (Figure 2D). Nevertheless, we investigated further the effects of ageing in each of the three tissues. In the ovaries, we detected 16 differentially targeted transposons, 7 of which were Penelope elements. The differential abundance was, however, marginal (Figure 1F). The piRNA profiles of thorax and brain were noisier, but we were able to define 11 and 7 differentially targeted transposons, respectively (Figure 1E&G). In the thorax, much like in the ovaries, these were mainly Penelope elements (6/11), whereas the transposons significantly affected in the brain mainly corresponded with lowly expressed piRNAs (Figure 1E). We conclude that piRNA expression changes only marginally with age in *T. rugatulus* queens.

**Figure 2:**
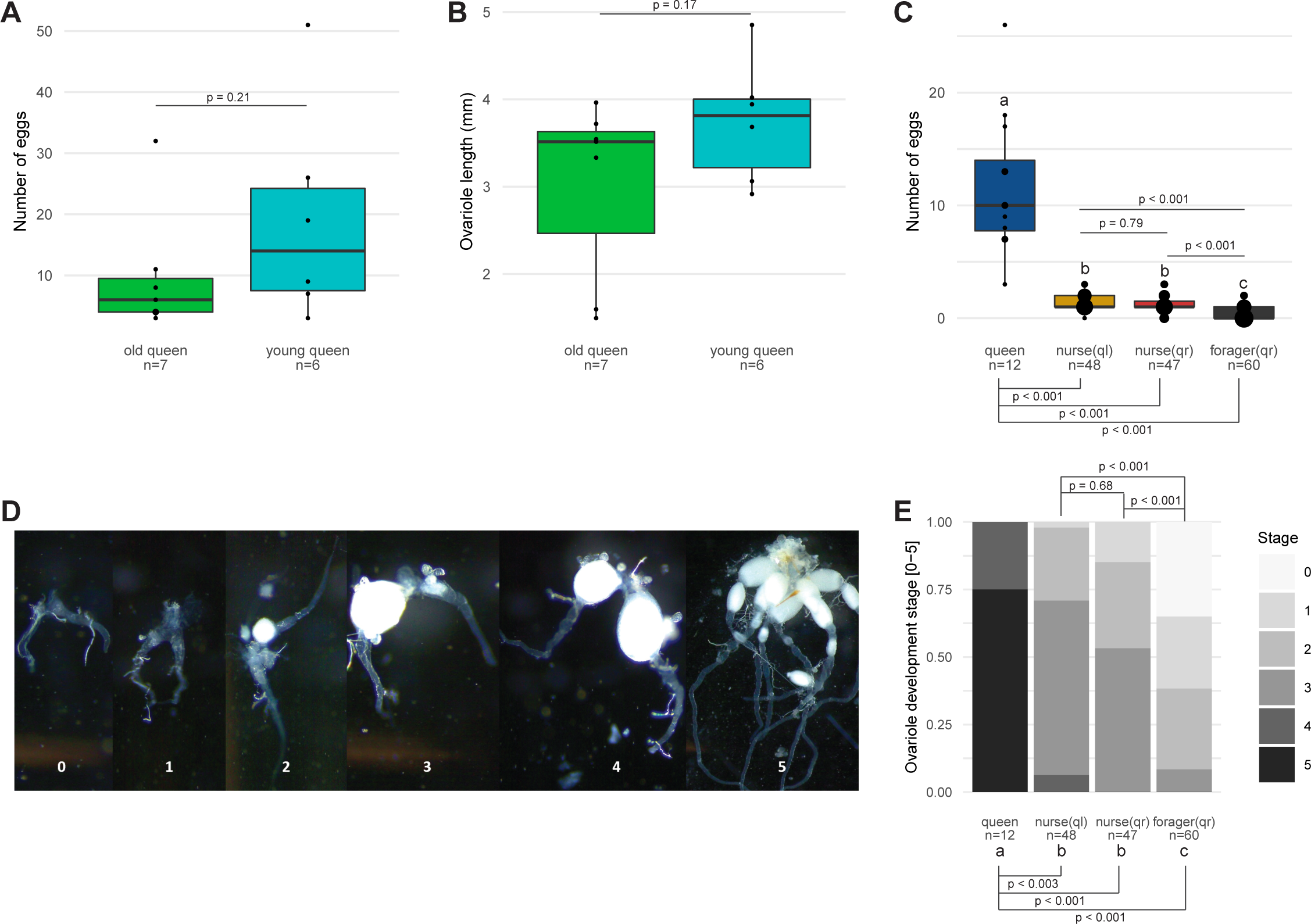
Eggs in development and ovarian development of different ages and castes of the ant *T. rugatulus*. **A** Boxplots of the number of eggs in development found in young, founding queens (< 6-month-old) and middle-aged established queens (> 3,5 years old). P-value for general linear model (quasipoisson) indicated. **B** Boxplot of ovariole lengths of young and old queens. Point size is relative to number of replicates. P-value for linear model indicated. **C** Boxplots of the number of eggs in development found in queens, nurses and foragers of qr colonies and nurse workers of ql colonies. Point size is relative to number of replicates. P-values for generalized linear mixed-effects model (family = poisson+ (1|Colony)) indicated, letiers indicate significance. **D** Representative pictures of five defined developmental stages of ovaries. 0 = regressed ovaries: thin, clear tubes; 1 = undeveloped ovaries with rounded tubes and a wider end, 2 = slightly developed ovaries containing immature eggs; 3 = developed ovaries with one mature egg, 4 = well developed ovaries with 2-5 mature eggs, 5 = extremely well-developed ovaries with > 5 eggs. E Boxplot of the distribution of each caste into the five developmental stages as defined in D. P-values for generalized linear mixedtieffects model (family = poisson) indicated, letiers indicate significance. qr = queenright, ql = queenless.

### Foragers have less developed ovaries than nurses and queens

Since we did not find differences in piRNA expression between old and young queens, we decided to determine their reproductive capabilities, thinking that low fertility and loss of ovarian piRNA expression might be associated. We dissected the ovaries of these seven old and six young *T. rugatulus* queens, measured ovariole length, and counted the number of developing eggs in their ovaries. The ovaries of old and young queens did not differ in the number of eggs in development (glm (quasipoisson): df = 1; χ² = 1.54; p = 0.21, Figure 2A), nor in ovariole length (glmer (family = poisson+ (1|Colony)): df = 3; χ² = 427.03; p < 2.2e-16); queen - ql-nurse: p < 0.001; queen – qr-forager: p < 0.001; queen – qr-nurse p < 0.001; qr-forager – ql-nurse p < 0.001; ql-nurse – qr-nurse p = 0.79; qr-nurse – qr-forager p < 0.001, Figure 2B). In contrast, we found strong differences in egg production and ovarian development between the different groups of queens and workers, when we dissected the ovaries of four different groups; queens, nurses, and foragers from normal colonies with a queen (queenright, qr), and nurses from colonies lacking a queen (queenless, ql). Queens had more eggs than all worker types, while foragers had fewer eggs in development than nurses (Figure 2C). We further defined the developmental stage of the ovaries ranging from 0 (undeveloped; no eggs in development) to 5 meaning (fully developed; more than five eggs in development) (Figure 2D). Here we could also see that queens had much more developed ovaries than any worker type whilst nurses generally had more developed ovaries than foragers (glmer (family = poisson): df = 3; χ² =72.483; p = 1.255e-15; queen - ql-nurse: p < 0.003; queen – qr-forager: p < 0.001; queen – qr-nurse p < 0.001; qr-forager – ql-nurse p < 0.001; ql-nurse – qr-nurse p = 0.68; qr-nurse – qr-forager p < 0.001, Figure 2E). We were surprised to observe no significant differences between eggs in development nor developmental stage between nurses from qr or ql colonies (Figure 2C&E), although those from ql colonies should have undergone ovarian activation in order to compensate for the lack of a queen and generally show a different gene activity (Negroni et al. 2021).

### piRNAs are expressed in the ovaries of queens and all worker types

After having determined that, in queens, piRNAs act primarily in the ovaries, we sought to investigate the piRNA profiles of ovaries from workers, even if they are only present as almost rudimentary organs. To allow an even comparison, we again sequenced sRNAs from the ovaries of queens, but this time also included the ovaries of foragers and nurses from qr colonies as well as nurses from ql colonies, where activation of the nurses’ ovaries are triggered. Analysing these four different groups allowed us to investigate differences in caste, worker task, and conditionary changes to nurses. Surprisingly, we obtained near-similar piRNA expression in the ovaries of all four groups (Figure 3A and Figure S2A) and only a slight depletion of ping-pong signatures in worker ovaries (Figure S2B).

**Figure 3:**
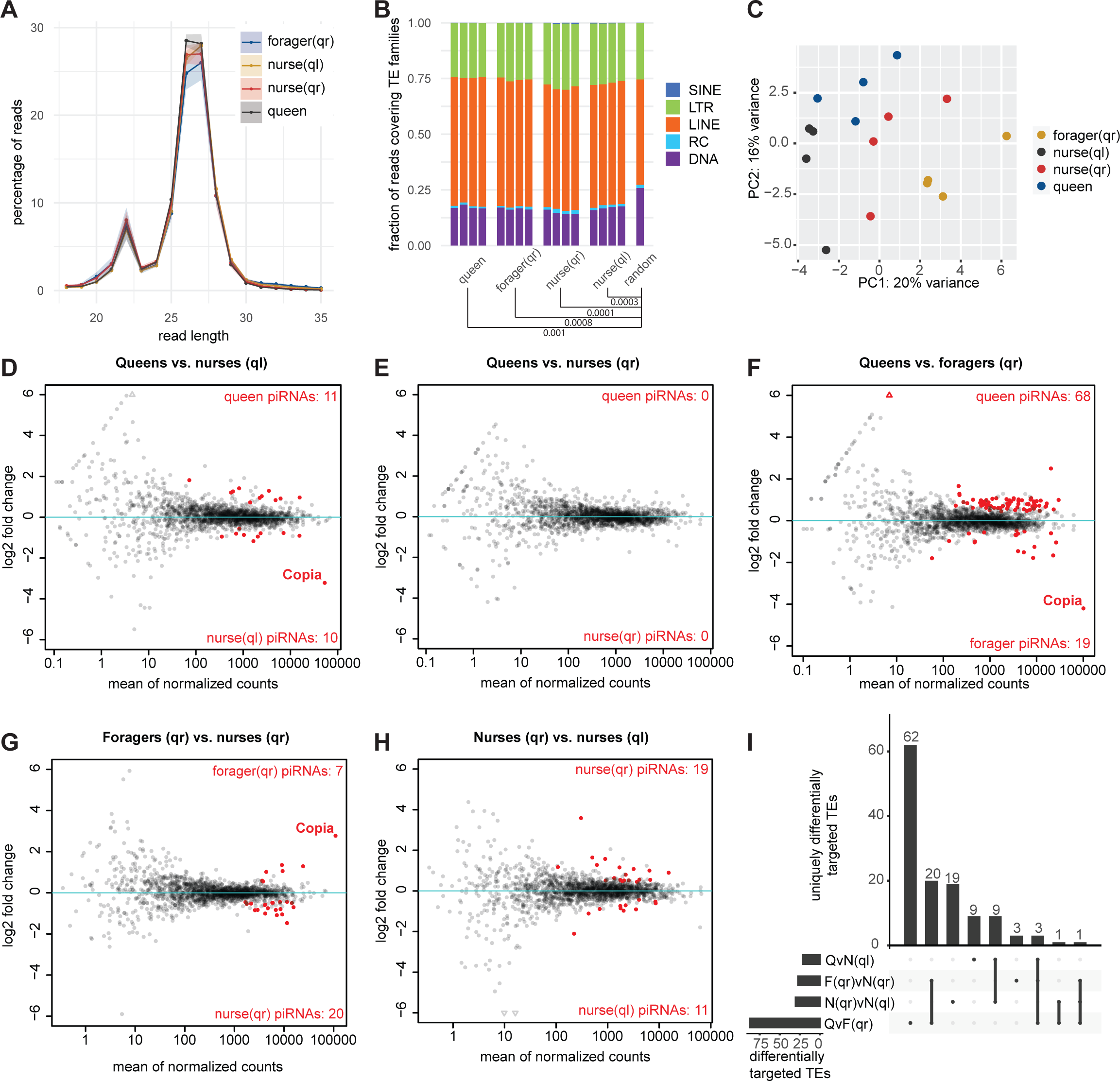
Ovarian piRNA expression in different castes. **A** Length distribution of all ovarian reads generated by sRNA-Seq. Shaded areas depict standard deviation, obtained from the 4 biological replicates. **B** Stacked bar plots showing piRNA-like reads atiributed to different TE families. T-tests carried out for each TE family to find differences between each caste and annotation. For each pair of comparisons, the lowest Holm-adjusted p-value is shown. All p-values listed in the supplementary data. **C** Principal component analysis of piRNA-like reads (25-30nt) that map to annotated TEs. **D-H** MA plots showing pairwise comparisons of piRNA targeting of specific TEs in the four castes. Each dot represents one TE. Red indicates significant changes (FDR<0.05). All hits are reported in the extended data, Table 2. **I** UpSet plot showing overlap of significantly differentially targeted TEs as defined in D-H. Hits per comparison are shown in the horizontal bar plot, overlaps between comparisons are 840 shown in the vertical bar plot. QvN(ql) = queens versus ql nurses, panel D. F(qr)vN(qr) = foragers versus qr nurses, panel G. N(qr)vN(ql) = qr nurses versus ql nurses, panel H. QvF(qr) = queens versus foragers, panel F.

As previously seen for the different queen tissues (Figure 1B), TE family targeting did not differ between these different groups either. Relative to the distribution of all annotated TEs, all samples showed a bias towards LINE elements at the expense of DNA elements (Figure 3B).

We carried out a differential analysis of our defined piRNAs using DESeq2 (Love et al., 2014) and observed only very small differences between the different groups, reflected in the minimal clustering in the principal component analysis (Figure 3C). Nonetheless, we could define several TEs with differential piRNA expression between groups with a particular upregulation of piRNAs in queens compared to foragers (Figure 3D-I). In general, those TEs that were found to be differentially targeted were those for which piRNAs were generally more expressed (Figure 3D-H), suggesting that, although the general upregulation of piRNAs in active versus inactive ovaries is modest (Figure 3A), the effect likely is biologically relevant.

We found one TE to be highly upregulated in foragers in several analyses (Figure 3D, F, and G), namely the Copia element. However, when further investigating the various Copia loci, we noticed that only one peak existed in each locus and that it consisted of only sense reads with no ping-pong signatures. We therefore extracted the sequences in the peaks and saw that these all aligned not only to each other, but also to the 3’ end of large subunit rRNA from *Bombus affjnis* (Figure S2C). Given the strong homologies of rRNA sequences between species, we conclude that Copia was not a real hit but rather an rRNA fragment that was relatively higher expressed in the foragers as a result of their slightly lower piRNA expression. This finding indicates that effects on individual TEs need to be treated with care. Therefore, we restricted our analyses to overall patterns and did not investigate individual loci.

We conclude that piRNAs are similarly active in the ovaries of all castes, with only marginal upregulation of already heavily targeted TEs in fully active ovaries. We also conclude that piRNAs are unlikely directly involved in the activation of ovaries in nurses upon removal of the queen.

### *T. rugatulus* express several miRNAs

We then investigated miRNA-like reads. First, we used miRDeep2 (Friedländer et al., 2012) to carry out a *de novo* miRNA prediction in our draft genome using miRNAs from four related species (*Bombyx mori*, *Drosophila melanogaster*, *Apis mellifera*, and *Tribolium castaneum*) as a reference and the pooled small RNA-seq data of all samples. Thereby, we could detect 1191 possible miRNAs. We then filtered these putative miRNAs using four parameters (see Methods) to ensure high fidelity of our generated miRNA annotation. After filtering, we were left with 372 miRNA loci coding for 304 distinct miRNAs (Table 3, extended data) Most of the miRNAs were unique, i.e. only found at one locus, and no miRNA was present at more than five loci. Notably, the identified miRNAs included strongly conserved miRNAs such as let-7 and mir-9, which are known to be ubiquitously expressed in other species.

After having generated our reference, we overlapped this with our mapped reads in the range 18-24nt from old and young queens. We saw that the reads mainly mapped sense to our defined miRNA loci, as would be expected for miRNAs (Figure 4A). We could further show that *T. rugatulus* miRNAs have a strong 5’-U bias (Figure 4A). A strong bias for 5’-U has also been indicated in other species, where this has been seen to aid in AGO recognition and direction into the correct AGO (Seitz et al., 2011). In *D. melanogaster*, for instance, a uridine at the 5’-positition will favour the uptake by the Argonaute AGO1 whereas a cytosine at the same position will favour uptake of the miRNA by AGO2 (Czech et al., 2009). In mammals, AGO1 preferentially binds 5’-U and AGO-2 preferentially binds 5’-A (Frank et al., 2010).

**Figure 4:**
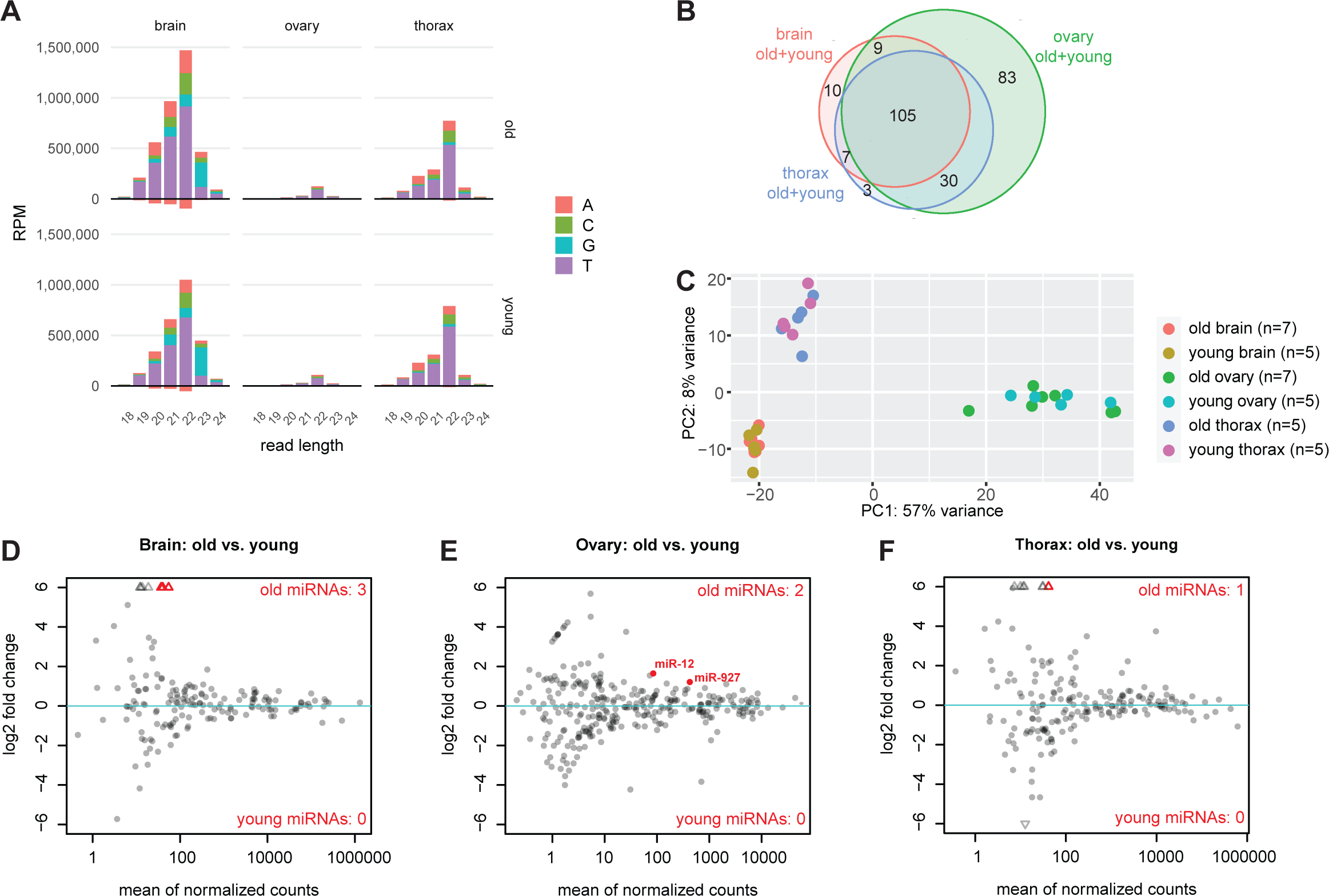
miRNA expression in different tissues of old and young queens. **A** Length distribution of reads length 18-24nt mapping sense (up) and antisense (down) to miRNAs. Values represent average of all 5-7 replicates, colours represent the starting base. **B** Venn diagram showing the overlap of distinct miRNAs found in at least three samples of brain, ovary, and thorax. Old and young queens are pooled together. All miRNAs are listed in the extended data, Table 4. **C** Principal component analysis of miRNA reads. **D-F** MA plots showing pairwise comparisons of miRNAs in old vs. young queens in the three tissues. Each dot represents one miRNA. Red indicates significant changes (FDR<0.05). All hits are reported in the extended data, Table 5.

Although relative miRNA expression was highest in the brain and thorax, ovaries expressed a more varied selection of miRNAs (Figure 4B). In the ovaries, we could detect the expression of 227 different miRNAs, 83 of which were unique for this tissue. In comparison, only 145 different miRNAs were detected in the thorax and 131 in the brain. We could detect 105 of the expressed miRNAs in all three tissues (Figure 4B). It is worth noting that expression here is defined as detection in at least three replicates of the same tissue, regardless of age, explaining why we find fewer expressed miRNAs (247 as shown in Figure 4B) than found in the genome (304, see above), where detection in three samples of *any* tissue was used as the filtering parameter. Altogether, we conclude that *T. rugatulus* does indeed express miRNAs and that these can be expressed tissue-specifically.

### miRNA expression is mostly not affected by queen age

We next carried out differential analyses in order to detect miRNAs involved in ageing. Similar to the piRNAs, the major difference in miRNA expression was related to tissue type, not age (Figure 4C). In the thorax and brain, we could define one and three miRNAs that were upregulated in the old queens, respectively. All of these were, however, lowly expressed (Figure 4D-F). In the ovaries, we found two miRNAs, miR-12 and miR-927a (both *A. mellifera* annotation), to be upregulated in the old queens, although both showed modest regulation (Figure 4E). In general, miRNAs seemed to be only mildly affected by ageing in *T. rugatulus* queens.

### Ovarian miRNA expression is specific to caste and worker task

Finally, we investigated the ovarian miRNAs of queens, nurses from qr and ql colonies, and foragers from qr colonies by isolating the reads in the range 18-24nt and mapping these to our miRNA annotation. Similar to results in queens only (Figure 4A), ovarian miRNAs of all groups mapped primarily sense to annotated miRNAs with a 5’-U bias (Figure 5A).

**Figure 5:**
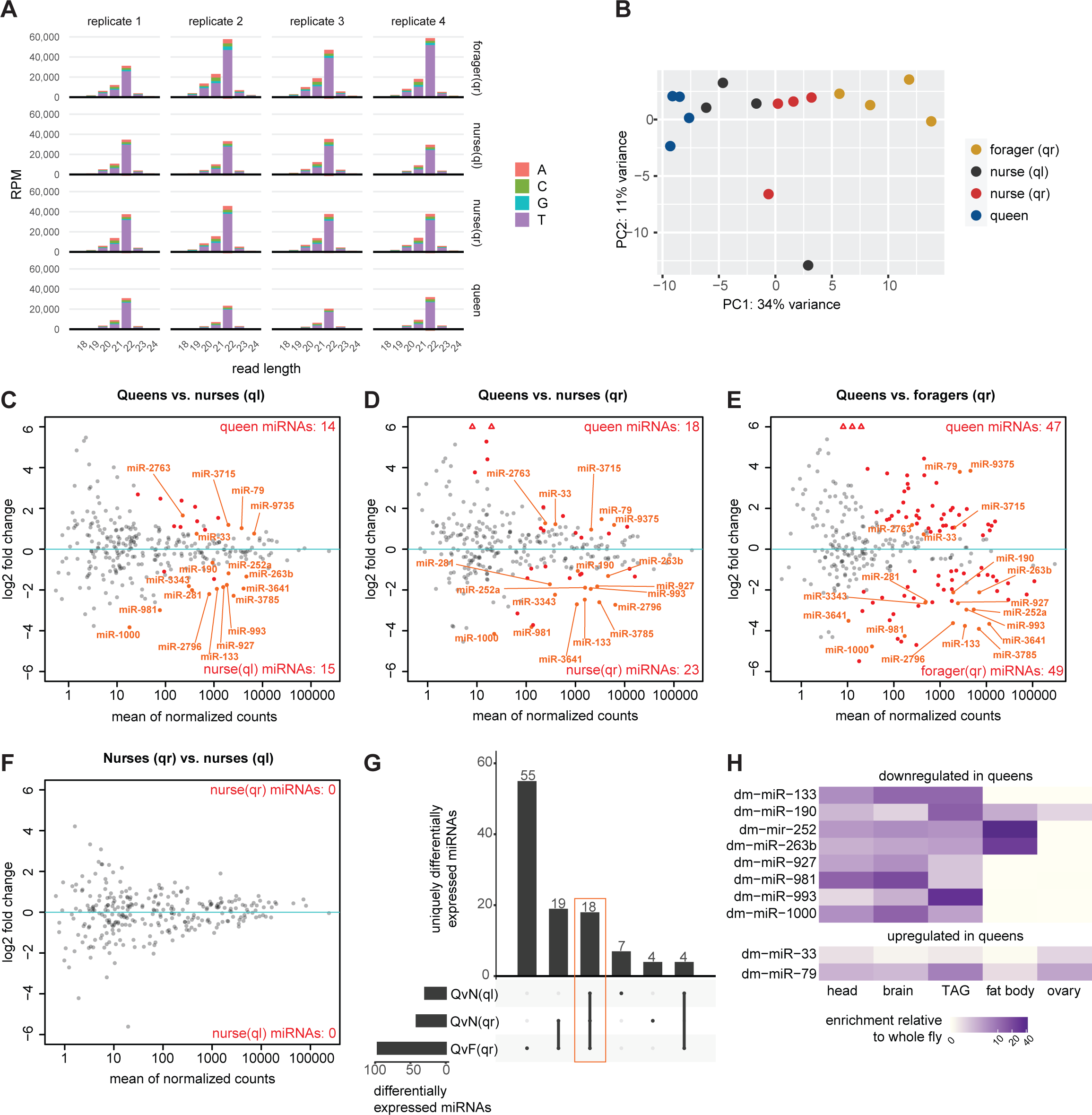
Ovarian miRNA expression in different castes. **A** Length distribution of ovarian reads length 18-24nt mapping to miRNAs on sense (up) and antisense (down) strands. Colours represent the starting base. **B** Principal component analysis of miRNA reads. **C-F** MA plots showing pairwise comparisons of miRNAs in different castes. Each dot represents one miRNA. Named miRNAs are those indicated in the orange square in panel G. Red indicates significant changes (FDR<0.05). All hits are reported in the extended data, Table 6. **G** UpSet plot showing overlap of significantly differentially expressed miRNAs as defined in C-F. Hits per comparison are shown in the horisontal bar plot, overlaps between comparisons are shown in the vertical bar plot. QvN(ql) = queens versus ql nurses, panel C. QvN(qr) = queens versus qr nurses, panel D. QvF(qr) = queens versus foragers, panel E. **H** Heatmap showing the relative expression of 12 miRNAs in D. melanogaster as reported in FlyAtlas2. These 12 miRNAs are those found in all comparisons C-E in five tissues (shown in the orange square in panel G) for which homologues exist in D. melanogaster. TAG = Thoracicoabdominal ganglion.

We next overlapped our mapped miRNAs with the miRNA annotation that we had already generated using our first dataset. In stark contrast to piRNAs, which showed little correlation between ovarian expression and group, we observed that miRNA expression and ovarian activation state were most likely associated. We observed a progression from active queen ovaries to activated ovaries of ql nurses onward to inactive ovaries of qr nurses and all the way to inactive ovaries of foragers, as visible in the principal component analysis (Figure 5B). Despite this segregation of our samples, we could not identify any specific miRNAs that could be directly involved in the activation of ovaries, as we did not detect any significantly regulated miRNAs between nurses of ql and qr colonies (Figure 5F). However, we could define more than 100 miRNAs that were differentially expressed in different castes across all comparisons made (Figure 5C-G, extended data Table 6).

We wanted to investigate a smaller subset of these miRNAs, and therefore decided to search FlyAtlas2 (Krause et al., 2022) for tissue-specific expression in *Drosophila melanogaster* of the 18 miRNAs that we found to be differentially expressed in queens in all three comparisons (queen vs ql nurses, queen vs qr nurses, and queen vs foragers). Two of the five miRNAs upregulated in queens and eight of the 13 miRNAs downregulated in queens had homologs with expression data in FlyAtlas2. We asked whether these miRNAs were generally known to be expressed in ovaries. Both of the queen-upregulated miRNAs are present in the *D. melanogaster* ovaries, but most of the queen-downregulated miRNAs did not show ovarian expression in *D. melanogaster* (Figure 5H). We noticed that all of the queen-downregulated miRNAs showed high expression in non-reproductive tissues; head, brain, and the thoracicoabdominal ganglion in *D. melanogaster* (Figure 5H). In other tissues such as the fat body, the expression level varied (Figure 5H). This indicates that non-queen ovary samples may carry significant amounts of other tissue(s). This could be due to their small size making them hard to dissect, or to the fact that these ovaries contain a lower fraction of germ cell mass, and hence relatively speaking more somatic tissue, or it could imply that these ovaries simply do express more miRNAs that are usually found in non-ovarian tissues.

Of note, we found miR-12 to be upregulated in the ovaries of queens relative to ql nurses (Figure 5C), and this coincides with the fact that we found miR-12 to be upregulated in old queen ovaries relative to young queen ovaries (Figure 4E). In the honeybee *A. mellifera*, miR-12 was found to be predominantly expressed in virgin and inactive ovaries (Macedo et al., 2016).

We conclude that ovarian miRNA expression is linked to caste identity and may be involved in the regulation of caste-specific genes.

## Discussion

### *T. rugatulus* expresses miRNAs and piRNAs

sRNA populations have been described in several fungi, plants, invertebrates, and mammals (Bartel, 2009; Ozata et al., 2019) and even analyzed in some ant species (Bonasio et al., 2010), but so far not described in *T. rugatulus*. We sampled *T. rugatulus* ants of different ages, castes, and worker types and sequenced sRNAs from brain, thorax and ovaries in order to describe the expression landscape of sRNAs across these different groups.

Our analyses confidently show that the ant *T. rugatulus* expresses two distinct sRNA populations coinciding with piRNAs and miRNAs, respectively. We also demonstrate that the relative expression of piRNAs in *T. rugatulus* queens is higher in the ovaries than in somatic tissues, which is consistent with the general function of piRNAs in reproductive tissues, where TE surveillance is particularly important (Madhani, 2013).

### Only minor changes of piRNA expression exist between different castes of *T. rugatulus* ants

Our results indicate that piRNAs may play a role in somatic tissues, most notably the thorax (Figure 1). This is consistent with a previous report describing piRNAs in somatic tissues in several ancestral arthropod species (Lewis et al., 2018).

There is growing evidence that sRNA pathways are linked to ageing, and increased TE activity has been a consistent molecular hallmark associated with ageing (Sturm, Perczel, Ivics, & Vellai, 2017). In the termite *M. bellicosus*, for example, genes associated with TEs were upregulated and genes from the piRNA pathway were downregulated in old workers with short residual lifespans, compared to long-lived reproductives (Elsner et al., 2018). On the contrary, TE expression did not increase in the fat body of the termite *Macrotermes natalensis* as the queens aged (Post et al., 2023). Furthermore, in *A. mellifera* TE activity and piRNA expression was only weakly correlated with both the highest level of piRNAs and TE activity in nonreproductive males (Wang et al., 2017), meaning that TE activity cannot necessarily be infered from piRNA levels alone.

We detected only 7-16 differentially targeted TEs between old and young queens (Figure 1E-G), all of which were only slightly regulated in the ovaries or showed only low sRNA levels in the brain and thorax. This suggests that sRNAs do not play a major role in the physiological ageing of this species. However, the small differences could also be due to the fact that we may not have included senescent queens at the end of their lives. We also note that the ants studied were field-collected rather than an established laboratory strain, and the fact that we detected any consistent changes in the first place may indicate that these indeed are true ageing-related differences.

In the termite *Macrotermes bellicosus*, TEs were found to be expressed in the heads of short-lived, old workers but not in queens (Elsner et al., 2018). This was also the case for the fat body of the termite *M. natalensis*, where TEs were found to be more active in workers than in reproductive individuals (Post et al., 2023), raising the question of whether the piRNA pathway targeting TEs would be less active in non-reproductive individuals compared to reproductive ones.

We demonstrate that the non-reproductive and regressed ovaries of qr nurses and foragers not only have piRNAs at similar levels to the queens, but they also show ping-pong signatures and target TEs which are all hallmarks of an active TE-silencing piRNA pathway. We note that piRNAs were slightly downregulated in foragers over queens (Figure 3A&F), but not nearly to the extent expected, given that the ovaries of foragers were regressed to the point that they were translucent membranes.

We conclude that piRNA populations change little upon ageing in *T. rugatulus*. They are most likely not the major driving force behind the ageing processes, with differences in other transcripts (Negroni et al., 2019), showing a much stronger correlation with queen age. In *T. rugatulus*, reproductive and non-reproductive ovaries alike seem to have the capacity to silence TEs via piRNAs and this pathway is likely not linked to ovarian activation. Rather, the high activity of piRNAs is a characteristic of ovarian tissue maintained even in the old workers, despite them sometimes being regarded as the soma of the superorganism ant colony (Johnson & Linksvayer, 2010).

### miRNA expression in relation to ageing and ovarian activation state

We detected 304 miRNAs in *T. rugatulus*, several of which are clear homologs of known miRNAs. Some of these are broadly expressed, but we also detected tissue specific expression. Many miRNAs have been shown to be differentially expressed during ageing across multiple organisms and this is associated with changes in the expression of genes involved in ageing and age-related diseases (Kinser & Pincus, 2020). This is for instance observed in the honeybee *A. mellifera*, where specific miRNAs were associated with age-dependent behavioral changes (Behura & Whitfield, 2010). We only found six differentially expressed miRNAs between old and young queens (Figure 4D-F), and these were either only slightly regulated or were expressed only at very low levels. Therefore, from our work, we cannot conclude that miRNAs play a role in ageing in *T. rugatulus*.

We detected larger differences in the miRNA populations between queens and foragers than between queens and nurses (Figure 5C-E). As nurses are younger than foragers (Kohlmeier et al., 2019), this could be partially attributed to an age difference, but since queen ovarian stage is more akin to that of nurses than that of foragers (Figure 2E), we propose that miRNAs more likely play a role in ovarian activity and possibly caste and/or task determination than in physiological ageing.

Our comparison of miRNA profiles of ovaries from *T. rugatulus* queens, foragers and nurses from qr colonies and nurses from ql colonies indicated a correlation between miRNA expression profiles and ovarian activation (Fig 5B). Several miRNAs were differentially expressed in individual comparisons between the reproductive ovaries of queens, and less reproductive ovaries of ql or qr nurses or foragers. We found no miRNAs to be differentially expressed between ql and qr nurses (Figure 5F), but also note that the differences between the number of eggs and ovarian development between these two groups were insignificant (Figure 2C&E). It has previously been shown that nurses of ql and qr colonies can differ in fat body gene expression linked and reproductive state (Negroni et al., 2019), but here we show that the link between miRNAs and ovarian activation is more subtle, and can only be detected when comparing all four groups (Figure 5B). One specific miRNA we detected in this context is miR-12, which we found to be upregulated in active ovaries of queens relative to ql nurses (Figure 5C). Interestingly, in *A. mellifera* (Macedo et al., 2016), miR-12, was found to be predominantly expressed in inactive ovaries, suggesting that miR-12 has a conserved role in ovary activation, but may have different target genes in *A. mellifera* compared to *T. rugutalus*. As more social insects have their miRNA profiles mapped, it will in the future become possible to do a more in-depth comparison and determine conserved versus species-specific caste-associated miRNAs and possibly to elucidate the evolutionary mechanisms behind the roles of sRNAs in the complex life styles of social insects.

## Acknowledgements

This project was funded by the Deutsche Forschungsgemeinschaft (DFG, German Research Foundation) – GRK2526/1 – Projectnr. 407023052. We thank the Genomics Core Facility at IMB for assistance in sequencing data.

## Data availability

Raw read libraries are available from the SRA database, BioProject PRJNA955004.

Draft genome and TE annotation are provided from Jongepier et al., 2021, available under BioProject PRJNA750352.

---

**Figure S1:**
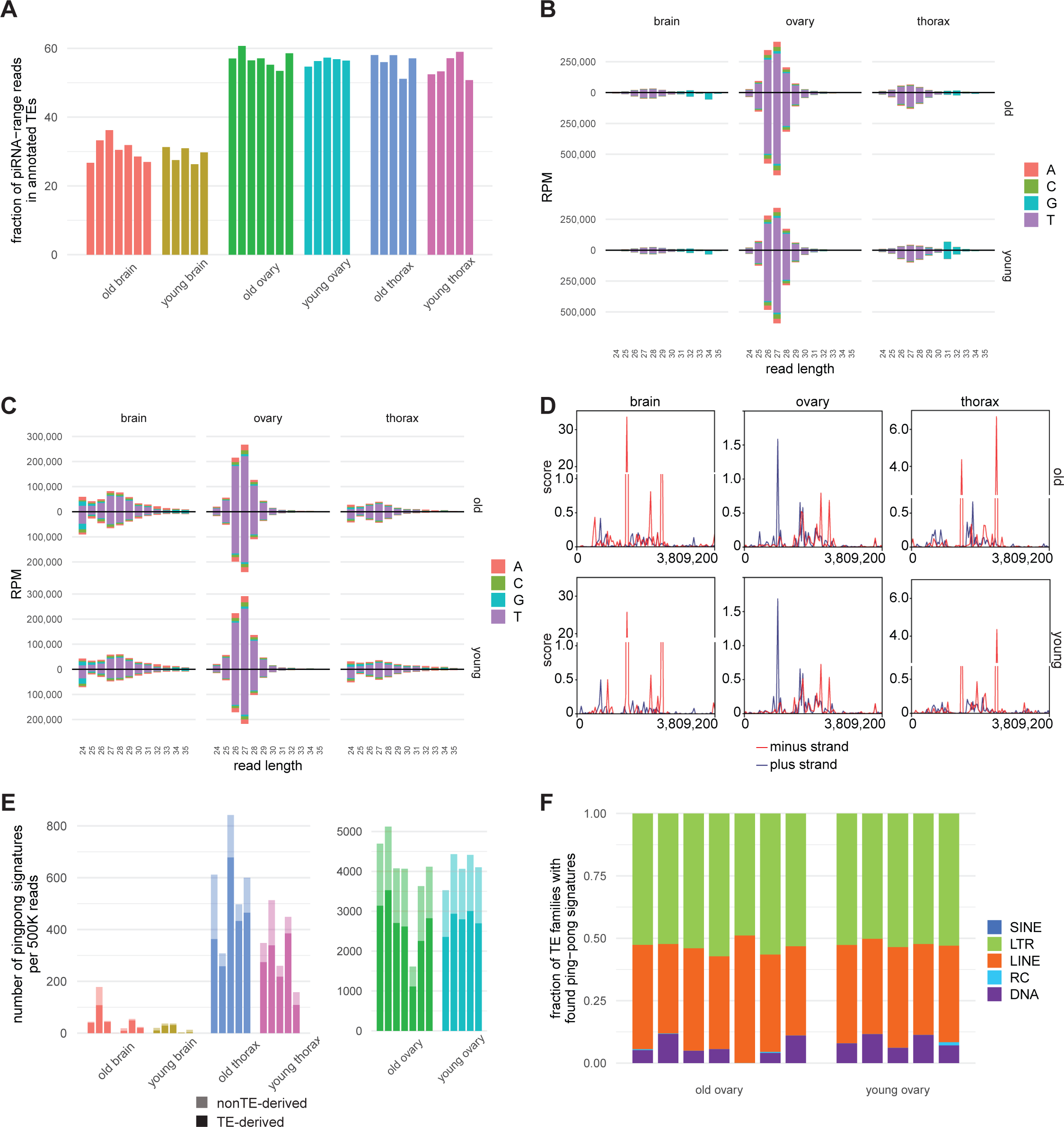
piRNA characteristics of 25-35nt sRNA reads in different tissues of queens. **A** Bar plot showing the fraction of piRNA-range reads (25-30nt) found within annotated TEs. **B** Length distribution of 24-35nt reads mapping sense (up) and antisense (down) to TEs. Colours represent the starting base. Reads were RPM normalized. **C** Length distribution of 24-35nt reads that do not map to annotated TEs. Reads are separated into plus strand (up) and minus strand (down). Colours represent the starting base. Reads were RPM normalized. **D** Coverage plot of piRNA-range reads mapping outside of TEs across the longest contig, 0031, which is 3.8Mbp in length, as indicated on the x-axis. Y-axis depicts the average read score of the 5-7 replicates on plus (blue) and minus (red) strands as calculated in bins of 10bp. **E** Bar plot showing the normalised number of ping-pong signatures found in each sample either within or outside of annotated TEs, as indicated below. Each replicate is depicted by a separate bar and replicates of the same age and tissue are shown in the same colour. T-tests old versus young: brain p = 0.23; thorax p = 0.07; ovary p = 0.66. **F** Stacked bar plot showing the fraction of TE-derived ping-pong pairs matching to each of the indicated TE families. T-tests old versus young: SINE p = 1; LTR p = 1; LINE p = 1; RC p = 1; DNA p = 1. T-tests old and young versus annotation: SINE p = 0.0; LTR p = 7.1e-13; LINE p = 4.3e-5; RC p = 3.8e-7; DNA p = 6.0e-9.

**Figure S2:**
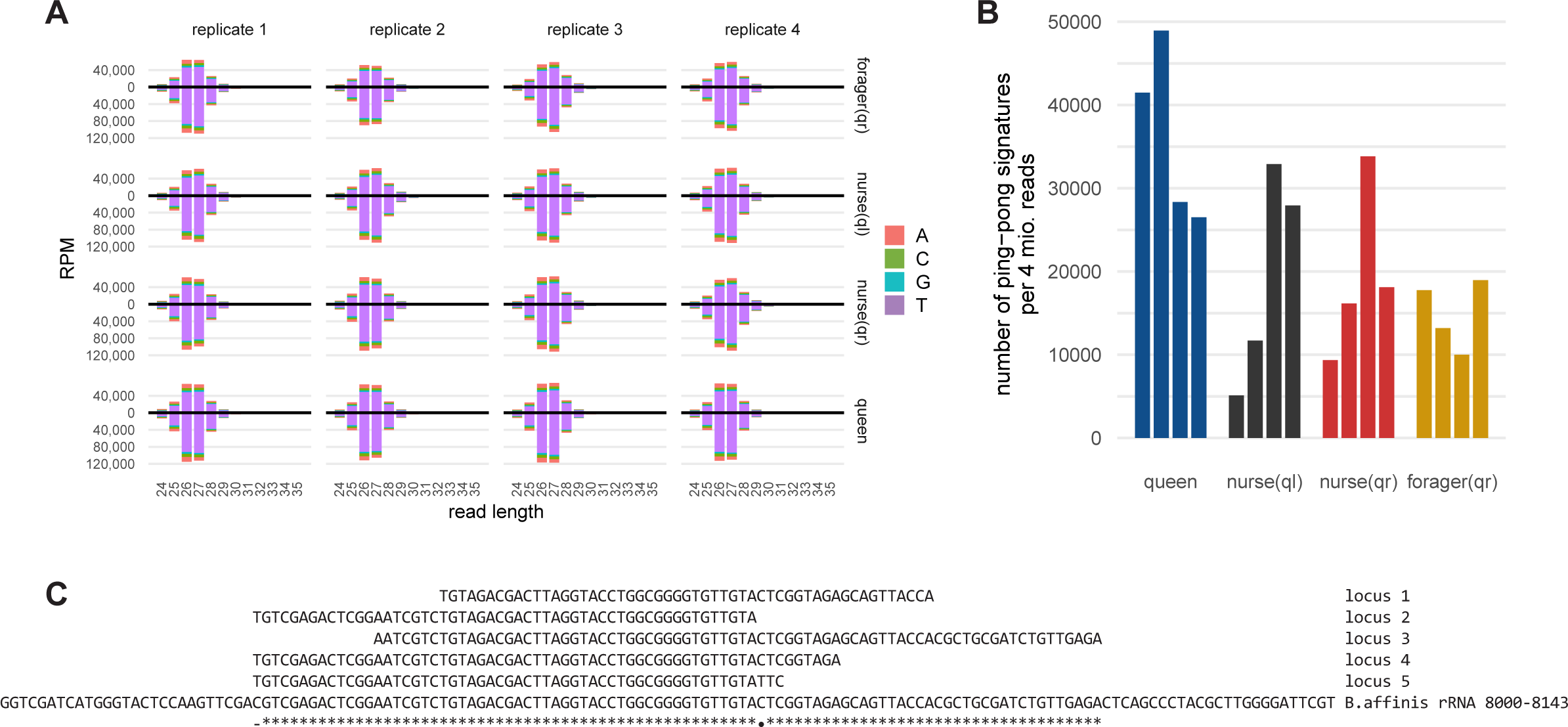
Ovarian piRNA characteristics of 25-35nt sRNA reads in different castes. **A** Length distribution of ovarian reads length 24-35nt mapping sense (up) and antisense (down) to TEs. Colours represent the starting base. **B** Bar plot showing the number of ping-pong signatures found in each sample. Replicates of the same caste are shown in the same colour. **C** Alignment of the sequences where the highest five peaks in Copia loci were found and the B. affinis large rRNA C-terminus, position 8000-8143. * 100% similar; • > 80% similar; - similar for loci, different from rRNA

## Notes

### Competing Interest Statement

The authors have declared no competing interest.

### Summary of Updates

An extra author has been added.

